# Real-time capture of horizontal gene transfers from gut microbiota by engineered CRISPR-Cas acquisition

**DOI:** 10.1101/492751

**Authors:** Christian Munck, Ravi U. Sheth, Daniel E. Freedberg, Harris H. Wang

## Abstract

Horizontal gene transfer (HGT) is central to the adaptation and evolution of bacteria. However, our knowledge about the flow of genetic material within complex microbiomes is lacking; most studies of HGT rely on bioinformatic analyses of genetic elements maintained on evolutionary timescales or experimental measurements of phenotypically trackable markers (e.g. antibiotic resistance). Consequently, our knowledge of the capacity and dynamics of HGT in complex communities is limited. Here, we utilize the CRISPR-Cas spacer acquisition process to detect HGT events from complex microbiota in real-time and at nucleotide resolution. In this system, a recording strain is exposed to a microbial sample, spacers are acquired from foreign transferred elements and permanently stored in genomic CRISPR arrays. Subsequently, sequencing and analysis of these spacers enables identification of the transferred elements. This approach allowed us to quantify transfer frequencies of individual mobile elements without the need for phenotypic markers or post-transfer replication. We show that HGT in human clinical fecal samples can be extensive and rapid, often involving multiple different plasmid types, with the IncX type being the most actively transferred. Importantly, the vast majority of transferred elements did not carry readily selectable phenotypic markers, highlighting the utility of our approach to reveal previously hidden real-time dynamics of mobile gene pools within complex microbiomes.

## Introduction

Densely populated polymicrobial communities exist ubiquitously in natural environments ranging from soil to the mammalian gastrointestinal tract. Bacteria in these microbiomes are thought to engage in extensive horizontal gene transfer (HGT) based on metagenomic sequencing studies and comparative genomics analyses^1–4^. HGT is a natural phenomenon where DNA is exchanged between organisms through distinct mechanisms including cell-to-cell conjugation of mobile plasmids or genetic elements, transduction by phages and viruses, or transformation by uptake of extracellular nucleic acids^5^. Upon horizontal transfer, the foreign genetic element can be either retained in the recipient or lost over time. HGT processes play a driving role in the evolution of bacterial genomes, leading to the dissemination of important functions such as complex carbohydrate metabolism^6^, pathogenicity^7^, and resistance to antibiotics^8^ or toxic compounds^9^.

Despite the prevalence of HGT, the evolutionary selection that drives fixation of foreign DNA is generally not well understood; for example, roughly 30% of genes predicted to be acquired by HGT have no known function^3^, and pan-genome analysis of sequenced genomes predict that many species have open-ended pan-genomes with enormous potential for gene turnover ^10–12^. For fixation of transferred DNA in recipient cells to ultimately occur, many barriers must be overcome, such as specific selection pressures, fitness burden of the acquired element, genetic compatibility with host machinery (e.g. replication, transcription, translation) and presence of anti-HGT systems such as restriction modification systems or CRISPR-Cas systems^5^. In addition, the presence of addiction elements on the transferred DNA (e.g. toxin-antitoxin and partitioning systems) also influence the fate of the transferred element. Even when the transferred genetic element provides a fitness benefit they may require many generations to be fixed in a population^13^. The architecture and dynamics of these gene flow networks are often not known, especially since most HGT genes are identified from endpoint analyses.

Contemporary computational methods for inference of HGT events rely on identification of shared mobile elements such as plasmids or phages, analysis of genomic abnormalities (e.g. shifts in GC% or codon usage) or phylogenetic comparisons between a candidate gene and a conserved gene (e.g. 16S rRNA)^14^. On the other hand, experimental approaches to study HGT require the transferred DNA to confer a detectable phenotype that can be enriched in the population. However, not all mobile elements confer a readily selectable phenotype. New selection-independent methods that can capture real-time transfer dynamics across a population will provide a deeper and richer understanding of the overall HGT process.

As a consequence of the pervasive gene flow in microbial genomes, bacteria have evolved various defense systems to manage horizontally acquired genetic material^5,15^. CRISPR-Cas systems can provide specific and adaptive immunity to invading DNA^15^. During the conserved CRISPR adaptation process, Cas1 and Cas2 proteins capture short fragments of the invading DNA and integrate them as spacers into CRISPR arrays. Immunity is conferred by transcribed spacers in conjunction with the Cas interference machinery^16^. Importantly, these CRISPR arrays provide a useful long-term record of horizontally acquired DNA. Most natural *E. coli* strains do not actively acquire new spacers and their arrays therefore reflect ancient HGT events^17^. However, overexpression of the Cas1 and Cas2 proteins can activate spacer acquisition in *E. coli* ^18^.

Here, we leverage the CRISPR spacer acquisition process as a mechanism for real-time recording of HGT events at nucleotide-resolution. Using an optimized acquisition system, we can capture transient HGT events and identify DNA transfers that cannot be easily detected with traditional methods. The performance and technical accuracy of this system was rigorously characterized using defined strains and communities. Application of the system to clinical human fecal samples revealed prevalent and diverse HGT events, shedding light on the dynamics and frequency of HGT in the mammalian gut microbiome.

## Results

### Exogenous HGT DNA can be identified using CRISPR spacer acquisition

We previously engineered a CRISPR-based temporal recording system that acquired new spacers from either endogenous genomic DNA or a copy-number inducible plasmid^19^. In this system, we utilized a “recording strain” (hereafter referred to as EcRec) consisting of the *E. coli* BL21 strain with the pRec-*ΔlacI* plasmid, which contains an anhydrotetracycline (aTc) inducible operon of the *E. coli* Type I-E *cas1* and *cas2* genes. Upon induction of the recording strain, over-expressed Cas1 and Cas2 proteins incorporate DNA protospacer sequences into a CRISPR array on the genome at high frequencies. Since *E. coli* BL21 lacks interference machinery, acquired spacers do not lead to CRISPR-mediated adaptive immunity^20^. The system can thus serve as a recorder of intracellular DNA. CRISPR expansions can be easily analyzed by PCR amplification of the CRISPR array from a population of recording cells, and, if needed, enrichment for arrays with new spacers can be achieved by a simple gel extraction of expanded array products. Subsequent deep amplicon sequencing can be used to assess the spacer repertoire^19^. While spacers can be acquired from both endogenous and exogenous DNA sources, including the genome, there is a strong preference to acquire spacers from high copy replicative plasmids and invading phages^21^. Given the capacity of the enhanced spacer acquisition system to record intracellular DNA at much higher efficiency than the wild-type system, we hypothesized that the system could be used as a sensitive method to reveal HGT events (**Fig. 1a**) that may only occur transiently or at a low-frequency across a cell population.

**Figure 1.**
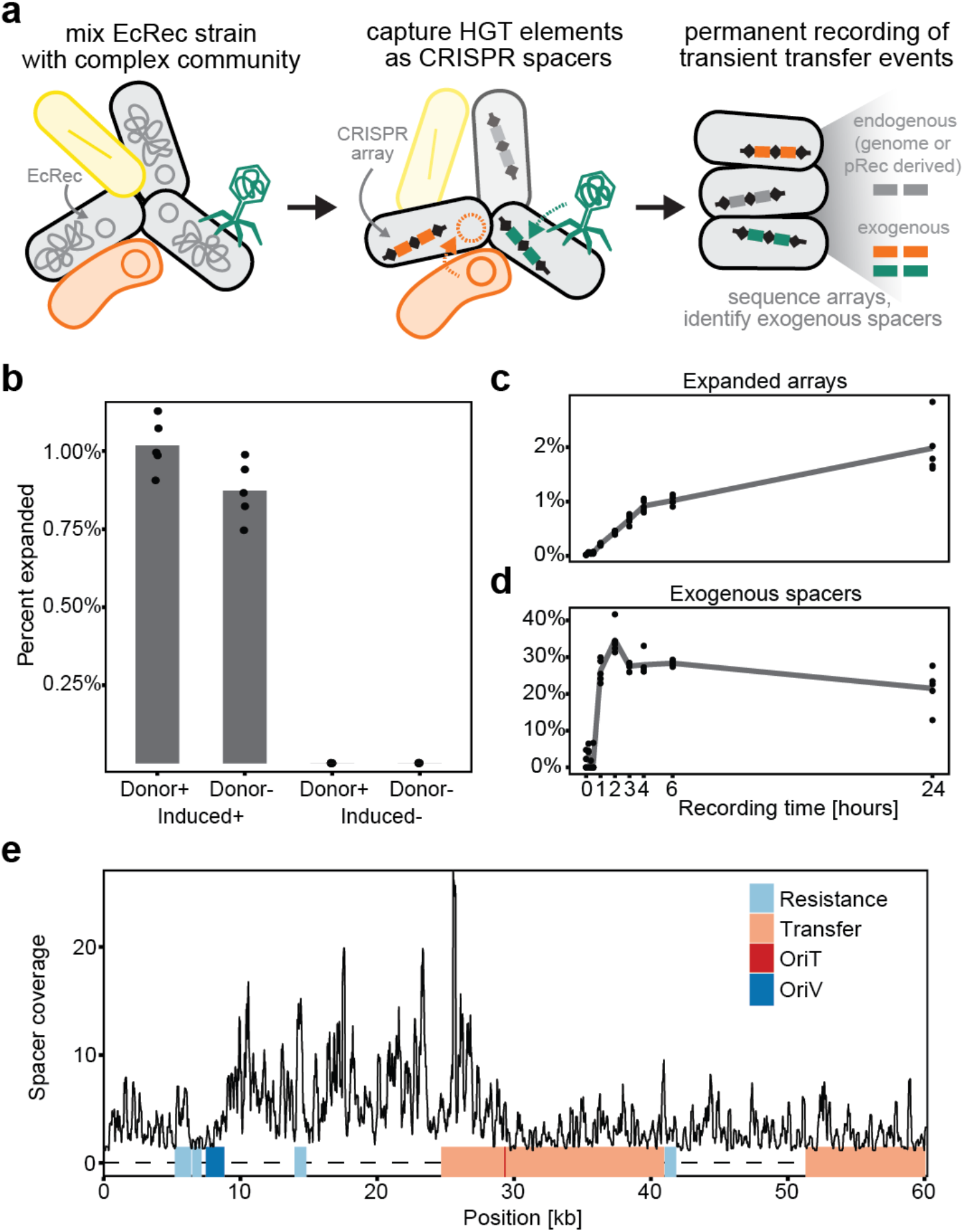
Recording HGT with engineered CRISPR acquisition. **(a)** Schematic of HGT recording where the EcRec strain is mixed with donor cells and spacers are acquired from both endogenous and exogenous DNA sources. Resulting CRISPR arrays are sequenced to determine the identity and origin of spacers. **(b)** Results from recording for 6 hours with or without induction and with or without FS1290/RP4 as donor strain (n = 5, with mean bar, no gel-extraction). **(c)** Array expansion is detected within 1 h after induction and increases rapidly for the first 4-6 hours (n = 5, with mean line, no gel-extraction). **(d)** Unique exogenous spacers are detected 1 h after induction, constituting ~30% of all spacers. **(e)** Mapping of recorded unique spacers to the RP4 plasmid. Spacer coverage is average coverage per bp. in 200 bp. windows and based on 5 replicates. On average, 99% of the unique exogenous spacers map to the RP4 and cover the whole plasmid backbone.

To explore whether CRISPR recording can allow direct measurement of HGT events, we exposed the recording strain (EcRec) to another strain (*E. coli* FS1290) that harbors the well-characterized broad host range conjugative plasmid RP4^22^. This conjugation was carried out by mixing the strains in a 1:1 ratio and spotting them on agar plates with and without induction of Cas1/Cas2. Reactions without the donor *E. coli* FS1290 strain served as an additional control. After 6 hours of conjugation, cells were collected and CRISPR arrays were amplified and sequenced (without gel extraction) to evaluate the spacer repertoire, yielding 10^4^-10^5^ sequenced arrays per biological replicate (**Suppl. Table 1**). In the Cas1/Cas2 induced cells with donor,1.0% (sd = 0.1%, n = 5 recordings) of the arrays were expanded in contrast to only 0.0010% (sd = 0.0006%, n = 5 recordings) in the non-induced cells (**Fig. 1b**). Further probing the dynamics of the HGT recording process showed that overall spacer expansion could be identified as early as 1 hour after mixing the donor and recording cells, with the rate of array expansion leveling off after 4 to 6 hours of bacterial conjugation (**Fig. 1c**). By 24 hours, 1.9% of all arrays (sd = 0.5%, n = 5 recordings) were expanded (**Suppl. Table 1**).

As expected, most spacers were derived from the EcRec genome and pRec plasmid. We therefore applied a stringent two-step filter against a *de novo* sequenced EcRec/pRec reference to isolate putative exogenous spacer sequences. First, only spacers flanked by the canonical direct repeat sequences were kept. Second, spacers with even moderate sequence homology (≥80% identity and coverage) to the EcRec genome or the pRec plasmid were removed (Materials and Methods and **Suppl. Fig S1**). Using these filtering criteria, we found that among the expanded arrays, exogenous spacers constituted up to 30-40% of all new spacers and could be detected within 1 hour of conjugation (**Fig. 1d**). After 24 hours, 21% (sd = 5%, n = 5 recordings) of the sequenced spacers were identified as exogenous. The amount of exogenous spacers is influenced by the ratio of donor to recording cells, and we could detect new exogenous spacers in as few as 1 donor per 10^6^ recording cells (**Suppl. Fig S2**). In comparison, only 0.5 % (sd = 0.2%, n = 5 recordings) of the spacers in the induced no-donor experiment were identified as exogenous, likely representing spacer sequences containing technical sequencing errors (**Suppl. Table 1**).

In complex microbiomes, the identity of potential transferred elements is unknown. However, acquired exogenous spacers can be matched against large sequence databases (e.g.GenBank) to identify specific mobile elements. To define the criteria for a match between a spacer and a reference database, we first gel extracted and sequenced spacers from the 24 hour *E. coli* FS1290 recording sample (**Suppl. Table 2**). A set of scrambled spacers was generated by randomly reordering the sequence of exogenous spacers. Using BLAST, both original and scrambled spacers were searched against the Genbank RefSeq bacterial genomes database. We identified a conservative threshold of ≥95% identity and coverage that prevented spurious matches of scrambled spacers to the database (**Suppl. Fig. S3**). Using this threshold, we found that 98.6% (sd = 0.2%, n = 5) of the unique exogenous spacers could be mapped back to the RP4 plasmid sequence (**Fig. 1e**) and that spacers were acquired across the plasmid, preferably from sites corresponding to the known PAM recognition sequence of *E. coli* Cas1/Cas2 (AAG, 50% of all spacers, **Suppl. Fig. S4**). Together, these results show that the EcRec is capable of recording horizontal gene transfer events robustly with high sensitivity and that exogenous spacers can be confidently mapped to the mobile DNA of origin.

### Detection of non-replicative and complex HGT events

Many HGT events may be transient and may occur at low frequencies. We hypothesized that our recording system could capture spacers from HGT events in which the transferred element is not stably maintained in a recipient. To investigate transfer of both genomic DNA and a non-replicative plasmid we used an *E. coli* S17 strain carrying the R6K-derived plasmid pUT, as the donor^23^. *E. coli* S17 contains a genomically integrated copy of the RP4-Tet::Mu conjugation system and also expresses the R6K replication initiation protein Pir. The integrated RP4 can mobilize the S17 genome and the pUT plasmid into recipient cells^24^. However, pUT requires the Pir protein *in trans* in order to replicate and therefore cannot be maintained in the EcRec recipient, which lacks the *pir* gene^23,25^. In addition, phage Mu, which is also present in S17, can be acquired by recipients either via conjugation of the S17 genome or via phage particles^25^.

We mixed EcRec with the *E. coli* S17/pUT donor strain and recorded spacers for 6 hours. Analysis of new exogenous spacers from the S17/pUT donor showed acquisition from both the integrated RP4-Tet::Mu and the pUT plasmid, highlighting that active replication of the transferred element is not required for spacer acquisition (**Fig. 2a**).

**Figure 2.**
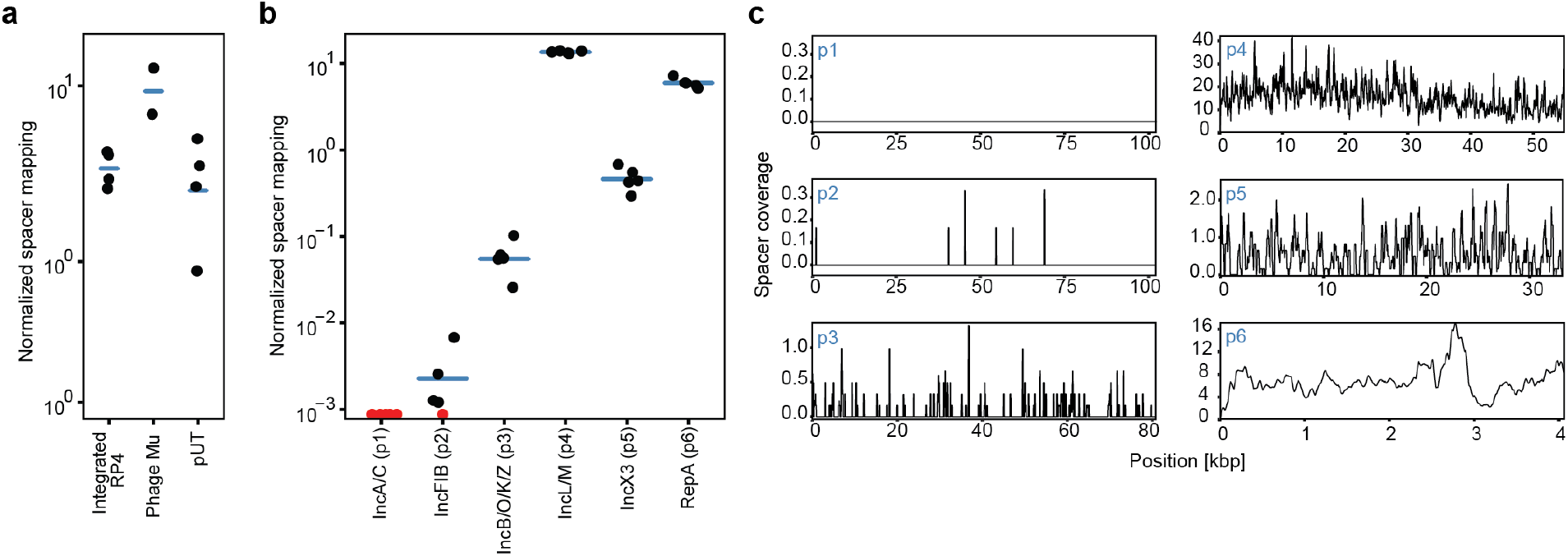
Detecting non-replicative and complex HGT events. **(a)** EcRec acquires spacers from transferrable but non-replicating DNA elements in *E. coli* S17, the integrated RP4, phage Mu and the nonreplicating plasmids pUT. In total 77,825 spacers were obtained. The normalized spacer mapping is spacers per kb per 1000 exogenous spacers. (n = 4 with mean bar, recorded for 6 h). **(b)** Recorded spacers from Ec70 carrying six plasmids (p1-p6). For each plasmid the number of matching spacers is normalized as spacers per kb per 1000 exogenous spacers. Red data points denote zero recorded spacers. No spacers are recorded from plasmid p1 (n = 5, with mean bar, recorded for 6 h). **(c)** Mapping of recorded spacers to the plasmid sequences, substantial coverage is seen for all plasmids except the large plasmids p1 and p2 (spacer coverage is average coverage per bp. in 200 bp. windows, based on 5 replicates.).

Since natural bacterial isolates often carry multiple plasmids capable of horizontal transfer, we tested if our HGT recording system could resolve transfer of different mobile elements from the same donor. A clinical *E. coli* isolate (Ec70) that carried 6 different plasmids (p1-p6), as resolved by hybrid assembly (Oxford Nanopore and Illumina sequencing, Materials and Methods), was used as the donor strain. Sequencing and analysis of new spacers from a recording experiment with Ec70 revealed that 97% of exogenous spacers were acquired from only two plasmids, the 55 kb plasmid p4 and the 4 kb plasmid p6. We quantified the spacer mapping to the reference sequence as the average number of spacers per kb per 1,000 sequenced exogenous spacers, hereafter referred to as “normalized spacer mapping” (**Fig. 2b**). For p4 and p6 the normalized spacer mapping was 13.5, sd = 0.4, and 6.0, sd = 0.8, respectively (n = 5). While plasmid p4 is self-transmissible, the much smaller plasmid p6 only carries the mobilization protein MobA, hence requiring the conjugation apparatus *in trans*. Neither of the two plasmids carry any antibiotic resistance genes, which importantly highlights that our recording system can readily detect elements that would not be easily detectable by standard selection-based methodologies. Plasmids p3 and p5 (80 kb and 33 kb, respectively) appeared to transfer, although at very low frequencies with a normalized spacer mapping of 0.060 and 0.48, respectively (sd = 0.03 and 0.15, n = 5 recordings). No spacers were observed from the large 106 kb plasmid p1 and only eight spacers were observed from the 102 kb plasmid p2 (**Fig. 2b**) from a total of ~1 million expanded spacers. As expected, spacers were acquired from across the plasmid backbones (**Fig. 2c**). Together, these results demonstrate that CRISPR-based recording of HGT can quantitatively reveal the relative transfer efficiencies of different mobile elements from a donor that carries 6 plasmids.

### Capturing HGT events from a defined microbial community

Having characterized the recording system using a single donor, we explored whether HGT events could be recorded in a complex, multi-donor community. A defined bacterial community comprised of 6 clinical *E. coli* isolates (Ec77, Ec70, Ec35, Ec14, Ec75, Ec21) as well as a positive control strain (FS1290) that carries the RP4 plasmid, and a negative control strain (REL606) that contains no plasmids was assembled. We generated draft genome assemblies and predicted that the clinical *E. coli* strains carried at least two plasmids each, including Ec70 already established to contain six plasmids^26^ (**Fig. 3**).

**Figure 3:**
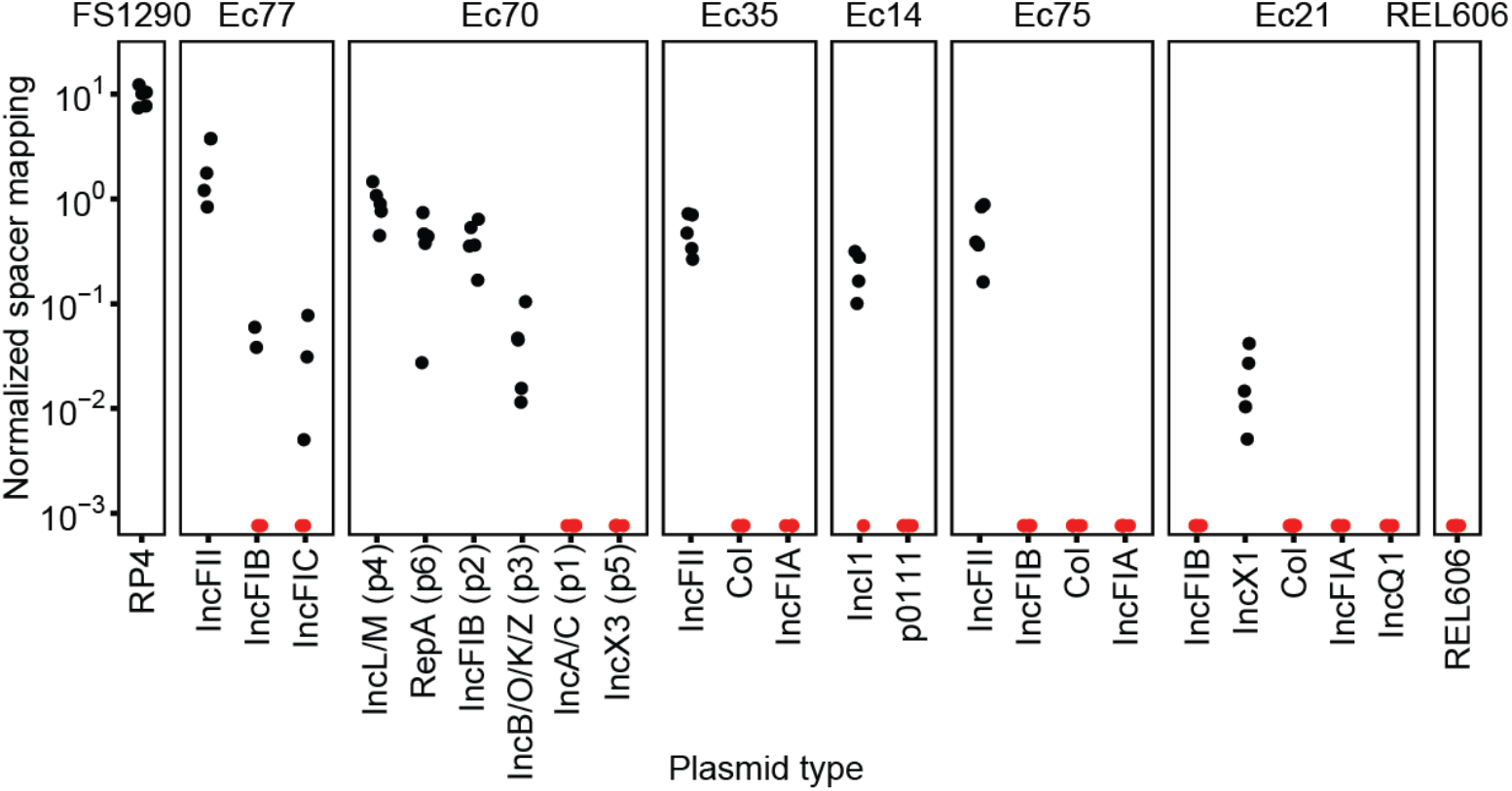
Recording of HGT events in a defined multispecies community. Spacer recording in a defined community of 8 *E. coli* strains. Exogenous spacers (n = 14,463 pooled over 5 biological replicates) were mapped to contigs identified as plasmids^26^ in the 8 genomes allowing only unique hits. Hits were observed for all donors except the negative control REL606, which carries no mobile genetic elements. The normalized spacer mapping is spacers per kb per 1000 exogenous spacers. Red data points denote zero recorded spacers.

**Figure 4.**
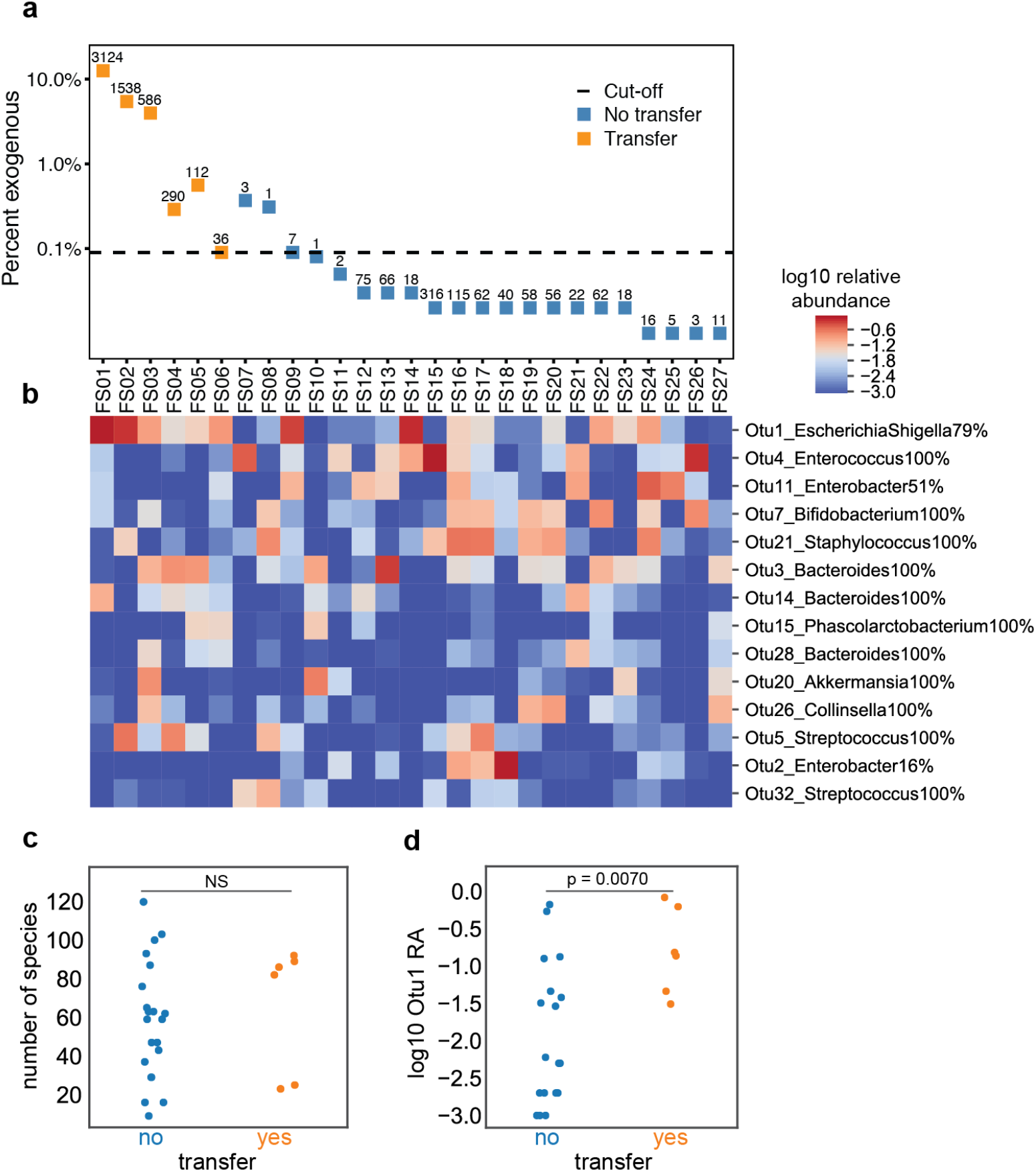
Measurement of HGT events in 27 clinical fecal samples. **(a)** Identifying fecal samples with robust exogenous spacer acquisition. Percent of spacers classified as exogenous for each sample. Samples with at least 0.09 % exogenous spacers and a minimum of 10 unique exogenous spacers (denoted above each data point) were classified as samples with HGT events (orange data points). **(b)** Cluster map of 16S OTU abundance for the 27 fecal samples. Samples with observable transfer (FS01- FS06) or no observable transfer (FS07-FS27) are shown. OTUs observed at >0.05 relative abundance in at least 2 samples are shown; log10 relative abundance is displayed. **(c)** Number of unique operational taxonomic units (OTUs) per samples stratified on transfer status. **(d)** Relative abundance of Otu1 (Escherichia/Shigella) stratified transfer status. Samples with transfer have a significant higher abundance of Escherichia/Shigella (p = 0.0070, Mann–Whitney U test).

Donor strains were pooled in equal ratios and then mixed with EcRec. Recording was carried out for 6 hours on LB agar and new exogenous spacers were identified and mapped back to the contigs from the draft genome assemblies for each of the 7 donor strains while the hybrid assembly was used for Ec70 strain. Spacers mapping to more than one contig were filtered out to ensure an unambiguous interpretation of HGT events (26.0%, n = 3205). We detected new spacers from all donor strains except from the negative control REL606 (**Fig. 3**). However, spacers were not acquired equally from the donors, with 72% (sd = 9%, n = 5 recordings) of all spacers deriving from the FS1290 positive control strain, confirming that RP4 transfers at high frequency^27^. Clinical strains Ec77 and Ec70 were particularly efficient donors, representing 19% and 7.3% of total spacers, respectively (sd = 9% and 2.4%, n = 5 recordings). Based on this mapping, we could also identify which predicted plasmids were or were not being transferred. For instance, IncFII-type plasmids in Ec35, Ec75, Ec77 appear to transfer readily to EcRec, while col-type plasmids in Ec21, Ec75, and Ec77 do not appear to mobilize. Importantly, we qualitatively detect the same transfer profile for Ec70 in this community recording as in the single donor recording. However, all spacers mapping to the IncX3 plasmid in Ec70 were removed due to redundant mapping to other plasmids in the community.

### Capturing HGT events from natural microbial communities

Extensive HGT has been reported in the human microbiome and has been shown to facilitate the spread of clinically important genes such as antibiotic resistance genes^3,4,28–30^. Therefore, we sought to explore the natural mobilome of the human fecal microbiome in clinically relevant populations using the CRISPR-recording system. Fecal samples were obtained from hospitalized adults with diarrhea whose stools were tested for *Clostridium difficile* infection (CDI). Of 27 patients samples, 24 had received broad-spectrum antibiotic treatment in the month prior to sampling while the remaining (FS05, FS06, and FS07) did not receive antibiotics. For each sample, ~0.5 g of fecal matter was washed in PBS three times to remove potential inhibiting compounds, such as antibiotics. The washed samples were each mixed with the EcRec, spotted on LB agar, and incubated for 24 hours. In order to confidently identify samples with HGT events we established strict criteria requiring that at least 10 exogenous spacers were identified and that the percentage of exogenous spacers was at least 3 times higher than in the no-donor control samples (0.03%). From the 27 recordings, we sequenced >10 million CRISPR arrays (**Fig. 5a**). Six recordings pass our criterion representing a total of 20,991 exogenous spacers yielding 5,686 unique spacers (**Suppl. Table 2**).

**Figure 5.**
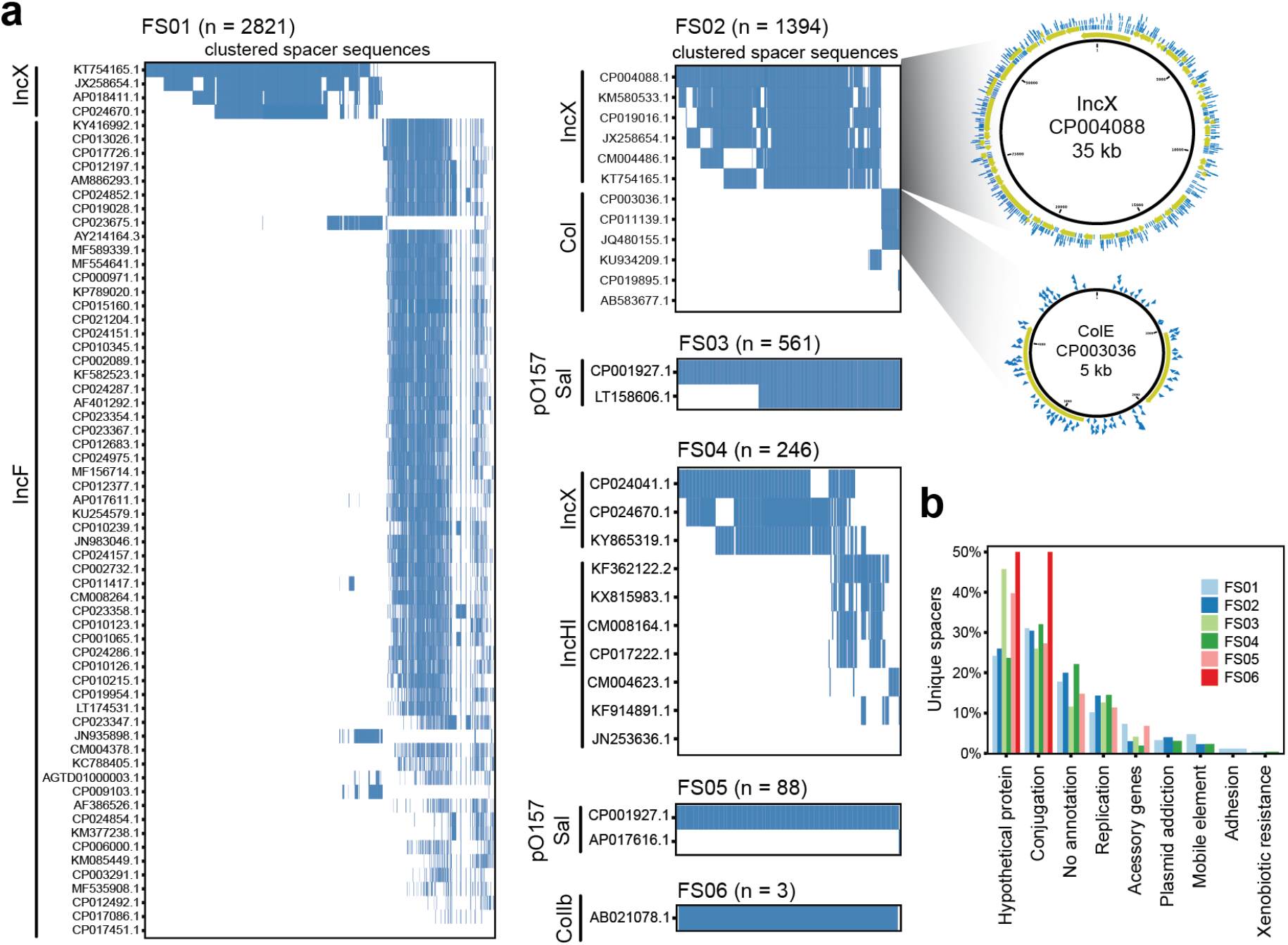
Analysis of human-associated mobilome from HGT recordings. **(a)** Exogenous spacers mapped to the custom plasmid database, each row represents a plasmid (denoted by accession number). The mappings are filtered to include the fewest number of plasmids covering all spacers. Rows are sorted in order of the number of spacers that map to the plasmid. The sorting enables easy identification of discrete transferred elements. Each spacer cluster is annotated with the predicted plasmid group based on Plasmid Finder^26^. Spacer mapping is illustrated for FS02 showing the plasmid backbone with predicted ORFs (yellow) and mapping spacers (blue). **(b)** Annotation categories overlapping with the spacers from all six clinical recordings. Genes predicted to be involved in conjugative transfer dominate, followed by unannotated genes and genes involved in plasmid replication. Notably, very few spacers overlap with genes involved in drug resistance.

We hypothesized that the presence of closely related donor species would be important to observing HGT. We thus profiled the composition of all 27 fecal samples using 16S rRNA amplicon sequencing (**Fig. 5b**). Overall, the a-diversity (number of species) was similar regardless of whether high numbers of exogenous spacers were observed (**Fig. 5b, c**). However, as predicted, we found that the relative abundance of the *Escherichia/Shigella* taxa (Otu1) was significantly elevated in the 6 samples passing the recording criteria (**Fig. 5d**, p = 0.007, Mann–Whitney U test). Still, some samples with high abundance of *Escherichia/Shigella* had few exogenous spacers (e.g. FS09 and FS14), suggesting that presence of *Escherichia/Shigella* at high abundance is correlated with but not sufficient for detectable transfer (e.g. presence and mobilization of plasmids may be variable in this host).

To identify the source of exogenous spacers, we used BLAST to search the NCBI RefSeq bacterial genome database, NCBI RefSeq viral genome database and a custom plasmid database applying the previously established thresholds (Materials and Methods).Overall, the majority of the 5,686 unique exogenous spacers could be matched to at least one of the databases (**Suppl. Table 2**). All spacers with hits to the viral database also matched to the genome database. Furthermore, >95% of spacers with hits to the genome database also matched to the plasmid database, highlighting that the identifiable spacers were most likely of plasmid origin. For each sample, we identified the minimal set of reference plasmids that encompass all spacers. Clustered heatmaps from these plasmid hits were used to identify the likely source of the exogenous spacers and predict the number of discrete mobile genetic elements. For each sample, we infer that 1-2 different plasmids were transferred (**Fig. 5a**).

For instance, BLAST hits of spacers to the plasmid database in sample FS02 indicate that two plasmids were transferred, a large IncX-type plasmid and a small colE-type plasmid (**Fig. 5a**). The putative IncX hits match to a 35 kb plasmid (Genbank accession CP004088) carrying no resistance markers. The acquired spacers almost completely tile the plasmid backbone, suggesting that the reference is a good representation of the transferred plasmid. The small colE-like plasmid (Genbank accession CP003036) has three predicted open reading frames (ORFs): a replication protein, a mobilization protein and an unknown ORF. While spacer coverage of the colE plasmid is sparser than the IncX plasmid, spacers matched across the back-bone suggesting that all regions of the pCE10B plasmid were present in the mobilized plasmid captured from FS02. Interestingly, the smaller plasmid does not encode a conjugation apparatus and therefore requires conjugation genes *in trans* for mobilization. Mapping all acquired spacers to the Plasmid Finder database^26^ revealed matches to IncX, IncI, IncF, IncH, pO157_Sal, and colE plasmid types, which are all common replicons in *Enterobacteriaceae* (**Fig. 5a**).

To better delineate the functions of the ORFs that yielded spacers, we used the RefSeq database to extract the functional annotations of genes with spacer hits (**Fig. 5b** and **Suppl. Table 4**). For each sample, 80-85% of spacers had functional annotations. The most common gene annotations were canonically associated with plasmids including conjugation, replication and plasmid addiction genes. As expected, a large portion of the ORFs had no known function (**Fig. 5b**). Given that the majority of patients received antibiotics recently (4/6 with detectable transfer), one might expect that these samples would be enriched in HGT for antibiotic resistance genes. Interestingly, mapping spacers against the ResFinder database^31^ yielded only two spacer hits to antibiotic resistance genes, a bla_TEM_ beta-lactamase and a chloramphenicol acetyltransferase gene (both from FS04), suggesting that, although present, resistance genes are not particularly enriched in these HGT pools.

### Systematic identification of transferred plasmids from metagenomes

We further performed shotgun metagenomic sequencing on the original fecal samples to assess the recovery of spacers against assembled contigs and to confirm the presence of putative plasmids in the samples. Metagenomic reads were assembled yielding ~371 Mbps of contigs across the six samples with recordable HGT events (FS01-FS06) (**Suppl. table 8**). Most acquired spacers could be matched to metagenomic contigs by BLAST (**Fig. 6a**). However, in two samples, FS05 and FS06, the metagenomic recovery rate was very low (3% and 8%, respectively). Correspondingly, these samples also had few acquired unique exogenous spacers (112 and 36, respectively), suggesting low frequency of HGT. Mapping of all exogenous spacers to the plasmid database revealed that the majority of spaces matched to both metagenomic contigs and published plasmids, confirming that most HGT was via plasmids (**Fig. 6a**).

**Figure 6.**
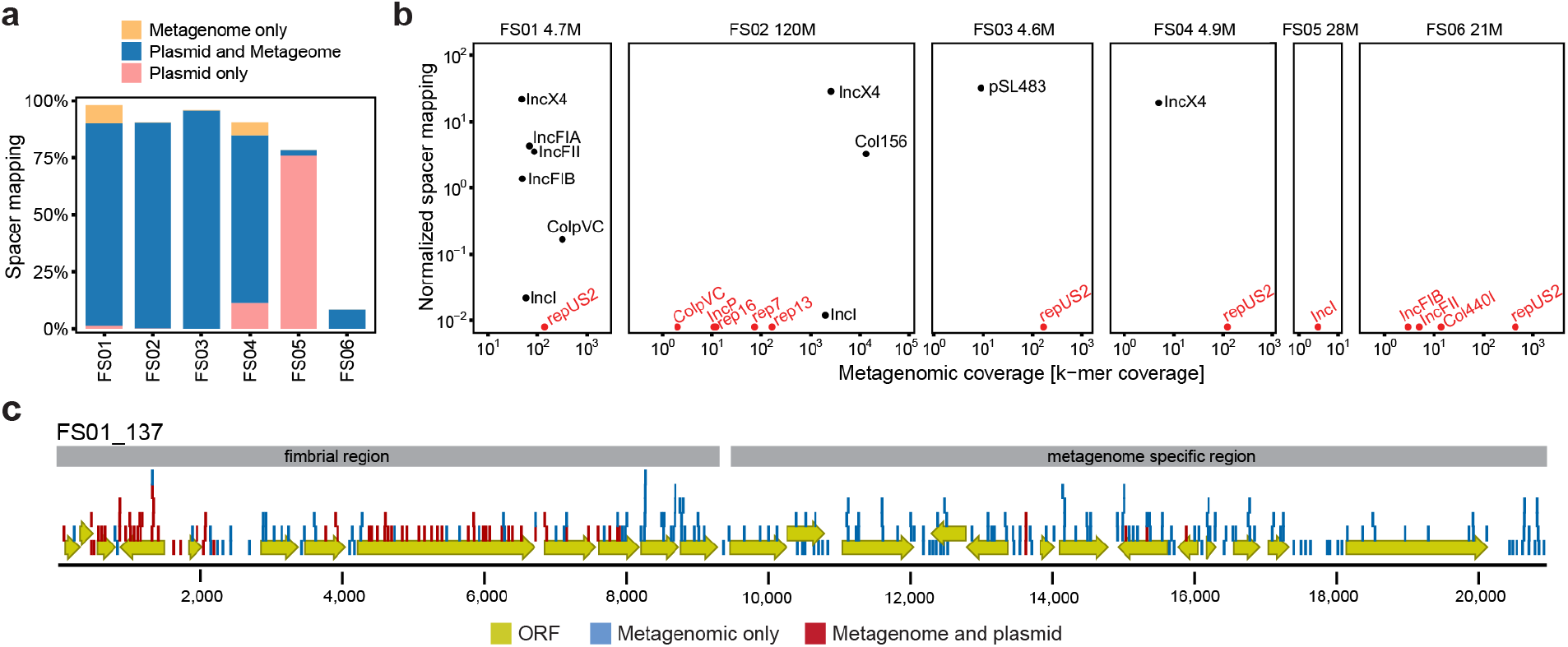
Metagenomic verification of predicted transfer events. **(a)** Percentage of spacers that could be mapped to the metagenomic contigs only (yellow), plasmid database and metagenomic contigs (blue), or plasmid database only (pink). **(b)** Mapping of spacers to predicted metagenomic plasmid contigs as a function of contig coverage in the assembly. The normalized spacer mapping is spacers per kb per 1000 exogenous spacers. Red data points denote zero recorded spacers. Number above each plot denotes the number of reads in the metagenome (millions). **(c)** Contig from FS01 where the majority of spacers were specific to the metagenome (blue). The contig consists of a region encoding a P-type fimbria and a region containing most hypothetical proteins specific to the metagenome.

Using the Plasmid Finder database^26^, we identified putative plasmid contigs across the metagenomes. We observed transfer of a variety of Enterobacteriaceae plasmids including IncF, IncX, IncI, and col types, corroborating the results generated from using our custom plasmid database (**Fig. 6b**). In addition, we also detected a number of non-transferred plasmids (e.g. repUS2) from Gram-positive species including S. *aureus*. Interestingly, certain plasmid types appeared to transfer more readily than others based on comparing their spacer mapping density and metagenomic coverage. In particular, IncX-type plasmids transferred efficiently since we observed nearly the same spacer density across three orders of magnitude in metagenomic coverage (**Fig. 6b**, FS01, FS02 and FS04). In contrast, IncI-type plasmids transferred at very low levels despite the metagenomic coverage varying two orders of magnitude between FS01 and FS02 (**Fig. 6b**).

A number of spacers mapped only to the metagenomic contigs (and not to the plasmid database; **Fig. 6a** and **Suppl. fig S5**). Among those contigs, one contig from FS01 had a majority of metagenomic-only spacers (202/276) indicating that the contig was not normally found on plasmids (**Fig. 6c**). This contig consists of a region encoding a P-type fimbria along with a transposase as well as a region containing mostly hypothetical proteins. The former region has been found in other plasmids, as indicated by spacer mapping to the plasmid database, while the latter region appears to be specific to the FS01 sample (**Fig. 6c**). The contig is not classified as plasmid, however, it is likely an incomplete assembly of a larger plasmid. This highlights the utility of our approach to identify novel transferred elements that may not be predicted by traditional reference-based methodologies.

### Estimating transfer frequencies in complex microbiomes

To assess the overall rate of HGT, including transient transfers, in our human fecal samples, we sequenced spacers from each fecal recording without gel extraction of the expanded arrays to calculate a rate of spacer acquisition (**Suppl. table 6**). The array expansion frequency ([# of expanded arrays] / [total # of arrays]) is multiplied by the frequency of unique exogenous spacers ([# of unique exo. spacers] / [# of expanded arrays]) to estimate the proportion of recording cells that captured exogenous spacers. For the six samples with high frequencies of exogenous spacers (FS01-FS06), the population-estimates of HGT spanned 4×10^-6^ to 10^-3^ [unique exo. spacers] / [total # of arrays] (**Suppl. table 7**). Together, these results suggest that while HGT may only be detected in some communities (e.g. 6 of 27 communities in this study), the extent of transfer in HGT-active communities can be quite high.

## Discussion

While comparative analyses of sequenced genomes have provided strong evidence of the abundant HGT^3,4^, the true rates of horizontal transfer of mobile DNA in a given community is poorly understood since many events may not fix in the population and diverse mobile elements compete for persistence in recipients. Our CRISPR-based recording system captures HGT events stably into genomic arrays that can be used to assess transfer rates and identity of mobile elements, far beyond current methodologies that rely on phenotypic selection of markers (e.g. co-transfer of antibiotic resistance genes). The ability to detect, in real-time, transient transfer events and those occurring at low frequencies enables an in-depth characterization of HGT in complex microbiomes. In this study, we showed that HGT can be resolved down to individual mobile plasmids from donors that can carry up to 6 putatively mobile plasmids. We find that the different microbial donors varied in transfer efficiencies of different plasmids, which might reflect differences in HGT efficiency between donor plasmids and/or competition for recipients. Such an approach could more generally facilitate detailed mechanistic studies of spread of mobile DNA associated with virulence phenotypes in specific pathogens.

When our approach is applied to clinical fecal specimens, we were able to identify active HGT in 22% of the samples (6 out of 27). In many instances, we observed multiple discrete plasmids being transferred, most of which interestingly did not carry selectable markers such as antibiotic resistance genes. This is surprising given the extensive usage of antibiotics in the cohort (24/27 patients). This finding suggests that a substantially larger pool of active and mobile plasmids exist in the gut microbiome beyond just the antibiotic resistance plasmids that are typically identified by phenotypic assays in experimental studies of HGT in the gut. By analyzing the captured spacers, we also find that many horizontally acquired genes have no known function, in agreement with previous bioinformatic analyses^3^. Using metagenomic sequencing, we definitively matched acquired spacer sequences to assembled plasmid contigs and plasmid types involved in these HGT events. While many different plasmids were identified in the metagenome, only subsets were shown to mobilize at varying efficiencies, with the IncX type transferring most efficiently.

The sensitivity of the spacer acquisition system allowed us to estimate the frequency of HGT in the human fecal samples. Because our estimates are based on adding a recipient strain to the fecal community, the transfer rates might not reflect actual transfer between community members, however, it does give an indication of the HGT potential of the community. We estimate transfer frequencies into the recording strain between 10^-6^-10^-3^ unique transferred spacers per recipient cell, suggesting that HGT is frequent.

Even though the observed mobile elements were classified plasmids, we still expect that phages are an important contributor to HGT. However, there are several possible explanations to why we do not observe phage-driven HGT. First, infection with phages could lead to cell lysis and consequently loss of recording cells from the population. Second, given that the *E. coli* CRISPR system is specific to capturing dsDNA, non-dsDNA phages (i.e. ssRNA or ssDNA phages) will not be captured. Third, the washing of the fecal sample (i.e. to remove antibiotics or other factors that might inhibit the recording strain) might result in the loss of most phage particles. We assessed the abundance of DNA phages relative to plasmids in the clinical fecal samples and found that on average there were 8 times more reads mapping to plasmids that to phages, suggesting that plasmids are more abundant in the fecal samples (**Supp. fig. S6**).

Future improvements to the technique could improve the scope of recording. Diverse CRISPR acquisition systems could be utilized to capture other HGT moieties (i.e. RNA with RT-Cas systems) beyond dsDNA captured by the *E. coli* system. Additionally, endogenous or engineered Cas1/Cas2 recording systems could be implemented in the context of different hosts to understand the host specificity of transfer for diverse bacterial species. These various systems and hosts could be multiplexed for high-resolution recording of HGT in various environments, from the human gut to various environmental microbiota. CRISPR spacer acquisition enables real-time recording of previously difficult to record transient HGT events, and offers a powerful new approach to studying flow and transfer of complex mobilomes at an unprecedented resolution.

## Supporting information

## Acknowledgements

We thank members of the Wang lab for helpful scientific discussions and feedback. H.H.W. acknowledges specific funding from ONR (N00014-17-1-2353), NSF (MCB-1453219), and NIH (1R01AI132403-01). C.M. acknowledges funding from the Carlsberg Foundation. R.U.S. is supported by a Fannie and John Hertz Foundation Fellowship and a NSF Graduate Research Fellowship (DGE-1644869). C.M. thanks Dr. Kristian Schønning, Hvidovre Hospital, Denmark, for gifting the clinical *E. coli* strains.

## Author contributions

C.M, R.U.S., and H.H.W. developed the initial concept. D.F. provided clinical samples and associated antibiotic treatment data. C.M, and R.U.S. performed experiments and analyzed the results under the supervision of H.H.W.; C.M., R.U.S. and H.H.W. wrote the mansucript with input from all authors.

## Materials and methods

### Strains

The recording strain (EcRec) was BL21 (NEB C2530H) with the pRec ΔlacI plasmid (Addgene #104575)^19^. Clinical *E. coli* isolates were a kind gift from Dr. Kristian Schønning, Hvidovre Hospital, Denmark. See **Suppl. table 5** for full overview of donor strains.

### Defined recordings

All strains were grown in LB medium with appropriate antibiotics and washed in PBS prior to recording. In all recordings an overnight culture of the recording strain was diluted 1:50 and grown for one hour, then anhydrotetracycline (aTc) was added to a final concentration of 100 ng / mL and the strain was incubated for another hour. Next, the recording strain and donor strain were mixed 1:1 at OD600 = 0.5, except in the ratio experiment (**Suppl. Fig. S1**) where strains were mixed in the ratios described in the figure. After mixing, the mixture was spotted on LB agar + 100 ng / mL aTc. Plates were incubated for 6 h at 37 C. At the end of a recording, the cells were scraped off the plate and resuspended in 100 μl PBS and heat inactivated at 95 C. for 3 min, subsequently they were stored at −20 C until sequencing analysis.

### Fecal recordings

The donor strain was prepared as described above. All fecal recordings were performed within 24 h of collecting the fecal samples. For each sample ~0.5 g were washed 2 times in 1 ml PBS and finally resuspended in 100 μl LB + 100 ng / ml aTc. The washed fecal sample was mixed with a 100 uL resuspension of 1 mL OD600 = 0.5 of the recording strain. From this mixture 50 μl was plated on LB agar + 100 ng / mL aTc and incubated for 24 h at 37 C. Subsequently, the samples were processed as described above.

### Ethical Review

The protocol for the collection of human samples and data was approved by the Columbia Institutional Review Board with a waiver of informed consent (IRB AAAR9489). Residual (waste) fecal specimens were used following standard clinical testing, and anonymized data was retrieved retrospectively.

### Array sequencing

CRISPR arrays were sequenced utilizing our established sequencing pipeline^19^ with minor modification. Briefly, DNA from cells was obtained by enzymatic and heat lysis, barcoded PCR amplification of CRISPR arrays was performed, samples were pooled and sequencing was performed on the Illumina MiSeq platform (MiSeq v2 50 cycle, MiSeq v2 300 cycle or MiSeq v3 150 cycle kits) with additional spike-in of custom sequencing primers. In addition, to enrich for expanded spacers, double gel extraction of expanded spacer bands on an E-gel EX Agarose Gel 2% was performed on pooled libraries. An overview of sequencing runs and sample statistics can be found in **Suppl. tables 1, 2, 6**

### Data processing

Spacers were extracted utilizing our established spacer extraction pipeline; code can be accessed at https://github.com/ravisheth/trace. Extracted spacers were filtered against the genome of the recording strain (quality filtered reads from sequencing of the same EcRec BL21/pRecΔlacI) using a two-step process using USEARCH v10.0.240^32^. First spacers were filtered using a database of word size 8, then all non-hit spacers were collected and filtered against the same database using word size 5 (e.g. ‘usearch -usearch_global -id 0.8 -query_cov0.8 -top_hit_only -maxrejects 0 -strand both -uc out.uc’). Subsequently the identified exogenous spacers were uniqued e.g.(‘usearch -fastx_uniques -fastaout centroids.fa -sizeout’). The unique exogenous spacers was utilized in all subsequent spacer mapping performed with BLAST 2.7.1+ (‘blastn -db -query -perc_identity 90 -max_target_seqs 500000000 -task blastn -word_size 10 -outfmt “6 std sstrand qlen slen”‘). The output of BLAST was filtered to ensure 95 % identity and 95 % coverage of the query spacer. An example of the processing workflow can be seen in **Suppl. Fig S1**. Data analysis was performed in R^33^ using ggplot2^34^ and CLC main workbench (www.qiagenbioinformatics.com).

### Reference databases

The following reference databases were used to identify the source of the acquired spacers: Prokaryotic RefSeq Genomes from January 2018; ftp://ftp.ncbi.nlm.nih.gov/genomes/refseq/bacteria/. Viral RefSeq Genomes from January 2018; ftp://ftp.ncbi.nlm.nih.gov/genomes/refseq/viral/. A custom plasmid database was created using the following search criteria in NCBI GenBank nucleotide database from January 2018; ‘plasmid[TI]’, then summary file was downloaded and parsed to get accession numbers of all circular elements: ‘grep -A1 ‘bp circular DNA’ summary.txt | grep -v ‘bp circular DNA’ | grep -v ‘\-\-’ | cut -d’ ‘-f1 > output.txt’ which were subsequently retrieved with NCBI batch (https://www.ncbi.nlm.nih.gov/sites/batchentrez).

### 16S rRNA sequencing

16S rRNA sequencing was performed utilizing our established sequencing pipeline; detailed methods can be found in our previous publication^35^. Briefly, genomic DNA (gDNA) was extracted with a protocol utilizing the Qiagen MagAttract PowerMicrobiome DNA/RNA kit (Qiagen 27500-4-EP). Samples were bead beat with 0.1mm Zirconia Silica Beads (Biospec 11079101Z) for a total of ten minutes (Biospec 1001); the Qiagen kit protocol was followed but at reduced volumes on a Biomek 4000 liquid handling robot. The resulting gDNA was subjected to 16S V4 amplicon sequencing utilizing custom barcoded primers^36^ and NEBNext Q5 Hot Start HiFi Master Mix (NEB M0543L). Resulting PCR products were quantified and pooled on a Biomek 4000 robot and sequenced utilizing an Illumina MiSeq V2 300 cycle kit. The sequencing data was analyzed using USEARCH 10.0.240^32^; reads were merged (-fastq_mergepairs), filtered (-fastq_filter -fastq_maxee 1.0 -fastq_minlen 240), and 100% ZOTUs were generated (-unoise3) and OTU table created (-otutab). Taxonomy was assigned to ZOTUs using the RDP classifier^37^. The OTU table was rarefied to 1000 reads per sample before analysis.

### Whole genome and shotgun metagenomic sequencing

The recording strain BL21/pRec along with all donor strains were subjected to whole genome sequencing (**Suppl. table 5**) and clinical samples were subjected to shotgun metagenomic sequencing (**Suppl. table 8**). gDNA was extracted from individual isolates or fecal samples utilizing the gDNA extraction pipeline detailed above. Sequencing preparation followed a published protocol for low-volume Nextera library preparation^38^. Barcoded samples were pooled and sequencing was performed on the Illumina MiSeq (2×150 reads), Illumina NextSeq (2×75 reads) or Illumina HiSeq X platform (2×150 reads). Adapters were trimmed utilizing Trimmomatic^39^. Draft assemblies for the donor strains were conducted using SPAdes utilizing the --careful flag^40^. Metagenomes were assembled with SPAdes utilizing the --meta flag. Raw metagenomic reads were mapped to the refseq viral database as well as the plasmid database using bwa mem^41^.

The donor strain Ec70 was further sequenced utilizing the Oxford MinION platform; genomic DNA was extracted with a Gentra Puregene kit (Qiagen), prepared for sequencing utilizing the RAD004 kit and sequenced on a single R9.4.1 flow cell. For this strain, hybrid assembly of the genome and individual plasmids was conducted utilizing UniCycler^42^. See **Suppl. table 5** for genome sequencing information and **Suppl. table 8** for metagenome sequencing information.

### Data availability

Assembled genomes, metagenomic reads, and CRISPR array sequencing will be made available at the time of publication through NCBI SRA.

