## Supplementary material for "Real-time capture of horizontal gene transfers from gut microbiota by engineered CRISPR-Cas acquisition"

**Supplementary Figure S1: Spacer analysis workflow.**
Spacers are first extracted and processed with the workflow below to first remove endogenous spacers and then match identified exogenous spacers to relevant databases.

1. **Extract spacers from raw sequencing files from Illumina instrument**

   Our previously published spacer extraction pipeline was utilized; code is available at https://github.com/ravisheth/trace

2. **Search spacers against the EcRec/pRec reference genome, using database with word size 8**

   ```
   usearch -usearch_global input.fa -db ref.reads.udb.fasta.8.udb -id 0.8 -query_cov 0.8 -
   top_hit_only -maxrejects 0 -strand both -uc out.uc
   ```

3. **Compile a fasta file with sequences not mapping to the word size 8 database**

   ```
   #Get the ids of the non hits
   find ./ -type f -name 'out.uc' | while read F
   do
     awk -F$'\t' '$1=="N" { print $9 }' ${F} > ${F}.exogenous.id.txt
   done
   #Compile a fasta file with the non hit sequences
   find ./ -type f -name 'input.fa' | while read F
   do
     grep -F -A1 -f ${F}.uc.exogenous.id.txt ${F} | sed '/^--/d' > ${F}.exogenous.ws8.fa
   done
   ```

4. **Search remaining spacers against the EcRec/pRec reference genome, using database with word size 5**

   ```
   usearch -usearch_global exogenous.ws8.fa -db ref.reads.udb.fasta.5.udb -id 0.8 -query_cov 0.8 -
   top_hit_only -maxrejects 0 -strand both -uc out.uc
   ```

5. **Compile a fasta file with sequences not mapping to word size 8 or word size 5 databases (i.e. exogenous spacers)**

   ```
   #Finally get all the exogenous spacers
   find ./ -type f -name 'out.uc' | while read F
   do
     awk -F$'\t' '$1=="N" { print $9 }' ${F} > ${F}.exogenous.id.txt
   done
   #Compile a fasta file with the non hit sequences
   find ./ -type f -name 'input.fa ' | while read F
   do
     grep -F -A1 -f ${F}.exogenous.ws8.fa.uc.exogenous.id.txt ${F} | sed '/^--/d' > ${F}.exogenous.fa
   done
   ```

6. **Cluster exogenous spacers**

   ```
   for file in *.exogenous.fa
   do
     usearch -fastx_uniques $file -fastaout $file.centroids.fa -sizeout
   done
   ```

### 7. BLAST unique exogenous spacers against desired database

```
blastn -db RefSeqJan2018 -query centroids.fa -perc_identity 90 -max_target_seqs 500000000 -task blastn -word_size 10 -num_threads 5 -outfmt "6 std sstrand qlen slen" -out centroids.refseq.hits.txt
```

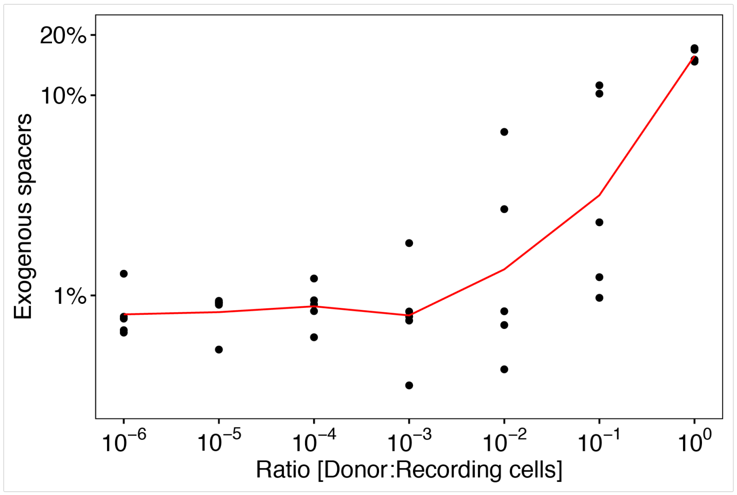


**Supplementary Figure S2: Effect of donor ration of spacer acquisition.**

Donor and EcRec was mixed in ratios from $10^{-6}$ – $10^{0}$ and spotted on LB agar. Recording was carried out for 6 hours.

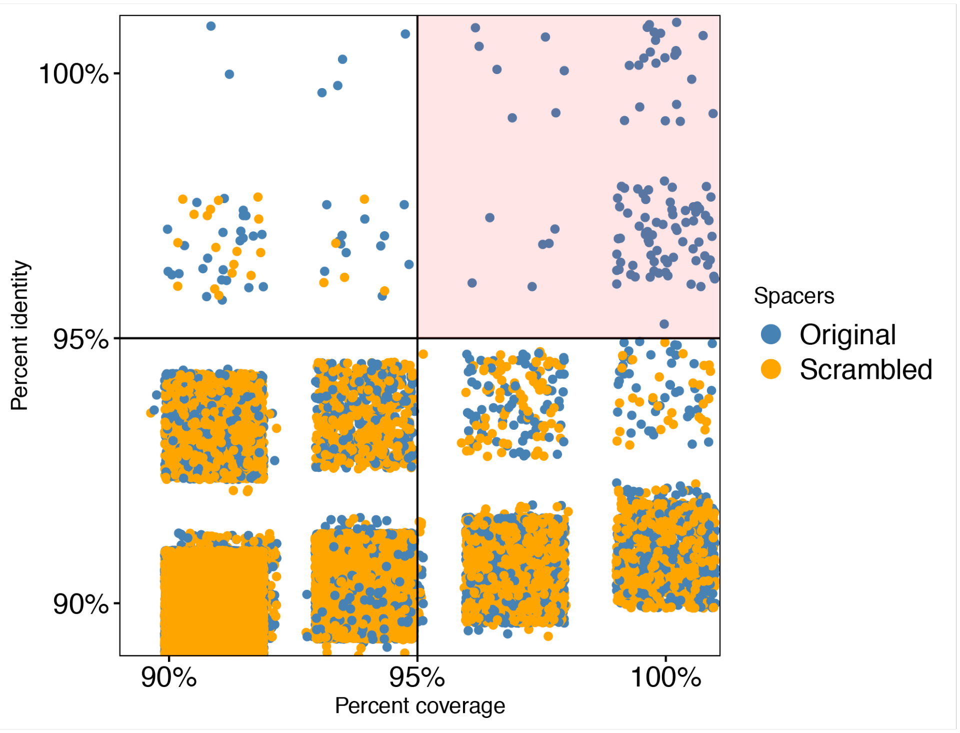


**Supplementary Figure S3: Identifying mapping cutoff.**
To identify cutoff for spacer mapping to databases of potential donors (e.g. Genbank nt) the recorded spacers from the E. coli FS1290/RP4 recording were scrambled by random reordering the sequence. Both the original and the scrambled spacers were mapped to the Genbank nt database using BLAST. We identified cutoffs of >=95% identity and coverage as resulting in reliable assignment of spacers (pink space). Each data point represent a unique spacer sequence.

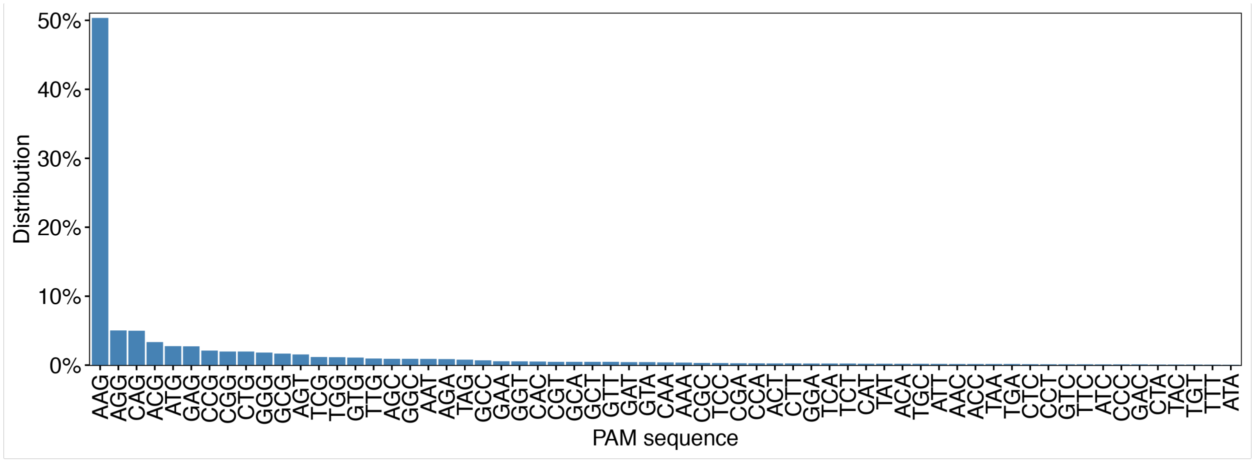


**Supplementary Figure S4: Distribution of protospacer adjacent motifs (PAM).**
PAM sequences were extracted for all spacers from the *E. coli* FS1290/RP4 mapping. The distribution shows a clear preference for spacers with the canonical AAG sequence.

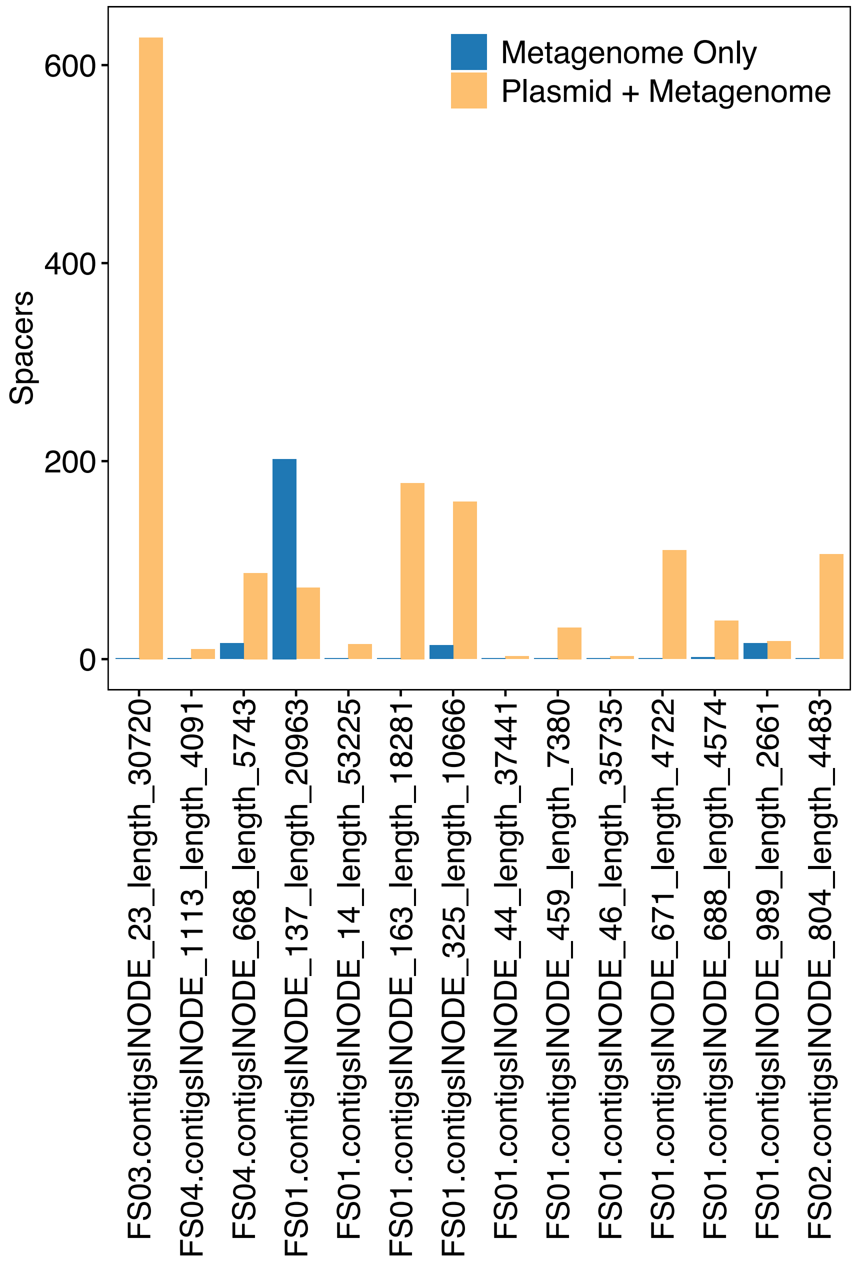


**Supplementary Figure S5: Contigs with metagenome-only spacers.**
Metagenomic contigs >500 bp that have at least two spacers mapping that do not map to the plasmid database. Shown is the number of spacers that only map to a metagenomic contig (blue bars) as well as spacers that map to both a metagenomic contig and a plasmid in the custom plasmid database. In all cases but one, most spacers mapping to a metagenomic contig also map to a plasmid contig indicating that the transferred element is known. However, in FS01 Node_137 the majority of the spacers only match to the metagenomic contig suggesting that most of this transferred element is not commonly found in plasmids.

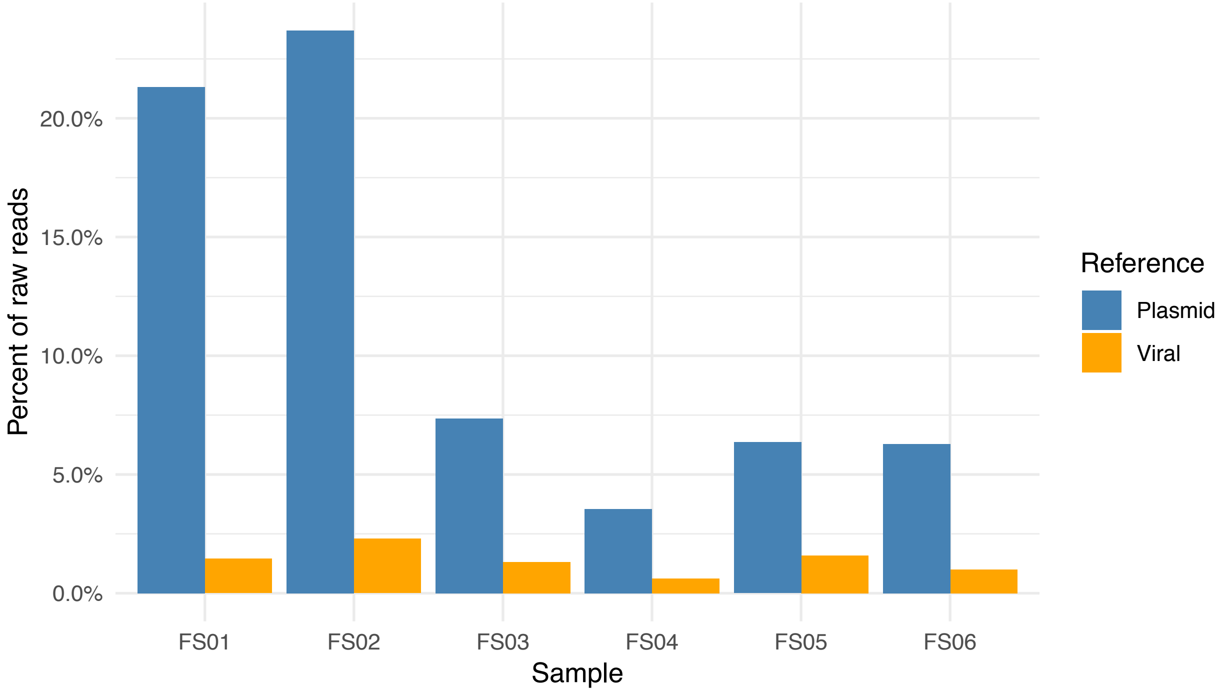


**Supplementary Figure S6: Metagenomic mapping to plasmid and viral databases.**
To investigate the abundance of plasmid reads vs. viral reads we mapped the raw metagenomic reads to the plasmid database and the refseq viral database, using bwa mem. Reads hitting to both databases were removed.
